# Cell-autonomous mitochondrial calcium flux governs oligodendrocyte regeneration

**DOI:** 10.1101/2025.11.18.689087

**Authors:** Dorien A. Maas, Roxane Bancel-Vega, Lucas R. Baudouin, Philippe Bun, Blandine Manot-Saillet, Chloé Habermacher, Niels Verburg, Filippo Rusconi, Maria Cecilia Angulo

## Abstract

Oligodendrocyte (OL) lineage cells drive central nervous system remyelination, yet the intrinsic mechanisms that define their regenerative potential remain unclear. We identify spontaneous, cell-autonomous intracellular Ca^2+^ signaling as a critical mechanism regulating OL regeneration following demyelination. Longitudinal *in vivo* imaging and *ex vivo* recordings reveal that Ca^2+^ signaling arises intrinsically within OL lineage cells after demyelination, and occurs independently from neuronal or behavioral activity. Mechanistically, mitochondrial Ca^2+^ flux sustains intracellular Ca^2+^ signals in oligodendroglia, and its *in vivo* disruption impairs oligodendrocyte precursor cell (OPC) proliferation, differentiation, and repopulation at the lesion site. Conversely, enhancing oligodendroglial Ca^2+^ signaling *in vivo* using chemogenetics stimulates lineage expansion and differentiation. In primary human OPC cultures, modulation of mitochondrial Ca^2+^ flux similarly reduces proliferation, indicating a conserved role for this pathway across species. These findings identify mitochondrial Ca^2+^ flux as a central driver of the oligodendroglial regeneration and a potential therapeutic target in demyelinating diseases.

## Introduction

Oligodendrocytes (OLs), the myelinating glia of the central nervous system (CNS), arise from oligodendrocyte precursor cells (OPCs) during development and in pathological conditions. In demyelinating diseases such as multiple sclerosis (MS), the loss of OLs denudes axons, compromising conduction and promoting neurodegeneration^1^. Although both surviving OLs and newly generated OPC-derived OLs can initiate remyelination^2–4^, this regenerative process is often incomplete or delayed, even at early disease stages. Unraveling the mechanisms underlying endogenous OL regeneration in demyelinated lesions remains an unmet challenge and a crucial first step toward developing novel remyelinating treatments.

OL regeneration in demyelinated lesions requires a coordinated sequence of cellular events, including OPC migration, proliferation and differentiation into myelinating OLs^1^. Although these cellular processes have been extensively characterized, the intrinsic signaling pathways that orchestrate the regenerative response remain unclear. Growing evidence suggests a role for intracellular Ca^2+^ signaling in oligodendroglia responses under both physiological and pathological conditions^5,6^. Intriguingly, Ca^2+^ signaling in OL declines with age, but is reactivated following remyelination^7^. However, it remains unknown whether this reactivation of Ca²⁺ signals simply reflects a consequence of the injury environment or instead represents a coordinated, active mechanism encompassing OPCs and OLs that governs regeneration after demyelination.

Like many cell types, oligodendroglial cells rely on intracellular Ca²⁺ as a universal second messenger to convert environmental cues into critical cellular responses such as proliferation and differentiation^5^. Recent *in vivo* studies have suggested that neuronal activity can modulate Ca^2+^ transients in the OL lineage. For instance, norepinephrine-mediated Ca²⁺ transients in OPCs increase during locomotion-induced arousal in mice^8,9^ and synaptically driven Ca²⁺ events in zebrafish OPCs occur at sites of future myelination^10^. However, whether neuronal activity is the primary driver of Ca^2+^ signaling in mature OLs remains debated: while one study showed that a substantial portion of Ca^2+^ transients in myelinating OLs are activity-dependent^11^, another reported predominantly activity-independent signaling in myelin sheaths^7^. This discrepancy raises a fundamental question: can oligodendroglia generate Ca²⁺ signals autonomously, independent of neuronal input? Furthermore, could these intrinsic signals be sufficient to govern OL regeneration?

Here, we demonstrate that spontaneous, cell-autonomous Ca^2+^ signals in OPCs and OLs appear in demyelinated lesions and are required for OL regeneration. Through longitudinal *in vivo* imaging, *ex vivo* slice recordings and targeted chemogenetic manipulations in transgenic mice, we show that Ca^2+^ activity in OPCs and OLs is intrinsically generated. Importantly, we identify mitochondrial Ca^2+^ flux as the key mechanism sustaining oligodendroglia Ca^2+^ signaling and regulating OL regeneration during remyelination. Disruption of mitochondrial Ca^2+^ flux impairs OL regenerative responses *in vivo*, consistent with its role in sustaining oligodendroglial Ca^2+^ signaling. Moreover, experiments in rodent and primary human OPC cultures indicate that this mitochondrial mechanism is required for initiating a proliferative process and that it is conserved across species, highlighting its potential relevance for human pathology. Altogether, these findings uncover a cell-autonomous, mitochondria-dependent Ca²⁺ signaling that governs oligodendroglia regeneration in the CNS, revealing an intrinsic organelle-based mechanism of glial repair that acts independently of neuronal input.

## Results

### Intrinsic Ca^2+^ signals emerge in OPCs and OLs within demyelinated lesions

While both OPCs and OLs display Ca^2+^ signals during development, how this signaling operates in demyelinated conditions and whether their dynamics depend on neuronal activity remains unclear. To investigate oligodendroglial Ca^2+^ signaling in demyelinated lesions, we employed the lysolecithin (LPC)-induced demyelination model in the corpus callosum (Fig. 1a). This model follows a well-defined sequence of events, with severe demyelination within the first 3 days post-injection (dpi), a peak of OPC proliferation around 7 dpi, and subsequent OPC differentiation and onset of remyelination between 7 dpi and 14 dpi, two critical stages for OL regeneration^12^.

**Figure 1.**
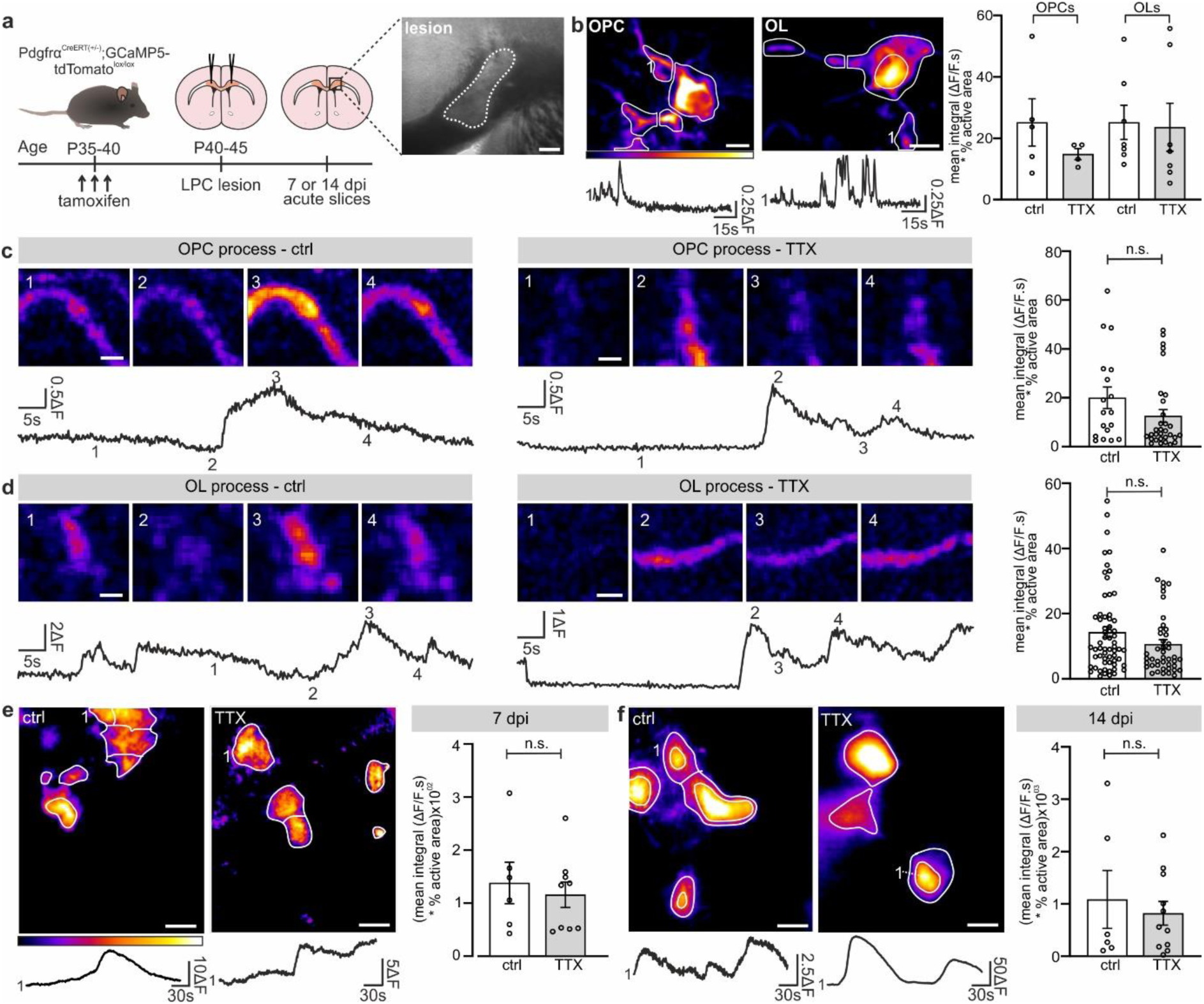
Oligodendroglial cells show spontaneous, intrinsic Ca^2+^ signals in lesions that do not depend on neuronal activity. (**a**) Experimental timeline. LPC-induced lesions (dotted line) were induced in *Pdgfrα^CreERT(+/-)^;Gcamp5-tdTomato^Lox/Lox^* mice expressing GCaMP5 in OL lineage cells and acute slices were prepared for Ca^2+^ imaging at 7dpi or 14dpi. (**b**) Representative images of two-photon microscopy recordings of an OPC and an OL expressing GCaMP5 (top) with designated active ROIs (white lines) as detected by Occam^6^. Representative Ca^2+^ traces are also shown (bottom). Dot plots (right) showing no differences in Ca^2+^ activity of OPCs and OLs between control conditions and in the presence of TTX (1 µM; n.s.: not significant; Kruskall-Wallis test followed by a post-hoc Dunn’s multiple comparisons test). (**c,d**) Representative images (top) and Ca^2+^ activity traces (bottom) recorded in processes of OPCs and OLs under control conditions and in the presence of TTX. Dot plots (right) showing no differences in Ca^2+^ activity in processes of OPCs (**c**) and OLs (**d**) under control conditions and in the presence of TTX (1µM; n.s.: not significant; two-tailed unpaired Student’s *t* test for OPCs, two-tailed Mann Whitney U test for OLs). (**e,f**) Representative projection image output by Occam with designated active ROIs (white lines) and Ca^2+^ activity traces (bottom) from widefield recordings in lesions under control conditions or in the presence of TTX. Ca^2+^ activity did not change under TTX exposure either at 7 dpi (**e**) or at 14 dpi (**f**). n.s.: not significant; two-tailed unpaired Student’s *t* test (**e**) and two-tailed Mann-Whitney U tests (**f**). Scale bars are 100 µm (**a**), 5 µm (**b**), 2 µm (**c,d**) and 20 µm (**e,f**). Data represented as mean ± standard error of the mean (s.e.m.).

First, we monitored Ca^2+^ dynamics in individual OPCs and OLs using conventional two-photon microscopy in acute brain slices from adult transgenic PdgfrαCre^ERT(+/-)^;Gcamp5-tdTomato^lox/lox^ mice (Fig. 1a). LPC-induced lesions were identified, at low magnification under the microscope, as brighter regions compared to the surrounding white matter, and single cells were then localized at high magnification (Fig. 1a,b). OPCs were distinguished by their relatively small, round somata and thin ramified processes, whereas OLs exhibited larger somata and prominent principal processes (Fig. 1b). As previously showed^6^, GCaMP5-expressing OPCs and OLs display complex spatiotemporal Ca^2+^ activity, making the detection of discrete events challenging. To capture this complexity, we quantified Ca^2+^ activity by calculating the mean intensity integral of automatically detected active regions of interest (ROIs), scaled by their active area, using the in house-developed open source programs Occam and post-prOccam^6^. We found that OPCs and OLs displayed comparable levels of activity in lesions, with most signals arising from their processes (Fig. 1b-d; see Supplementary Video 1).

To determine whether this Ca^2+^ activity was intrinsically or neuronally elicited, we blocked neuronal activity with the Na^+^ channel blocker tetrodotoxin (TTX) in acute slices. Notably, we found that similar levels of Ca^2+^ activity persisted in both somata and processes of OPCs and OLs in the presence of TTX (Fig. 1b-d), indicating that the observed Ca^2+^ signals in lesions are predominantly intrinsic to these glial cells, rather than elicited by neuronal activity.

To assess Ca^2+^ activity across larger lesion areas and compare the oligodendroglial population at 7 dpi and 14 dpi, we performed widefield imaging in the same transgenic mouse line. Although this approach does not allow resolution of individual cells, we observed that the overall population Ca²⁺ activity was comparable at both time points: 7 dpi when OPCs predominate, and 14 dpi, when OLs dominate the lesion (Fig. 1e,f; Supplementary Video 2)^12^. Moreover, Ca^2+^ activity remained insensitive to TTX, further confirming that oligodendroglial Ca^2+^ signals in lesions are intrinsically generated. To validate these findings with the more sensitive Ca^2+^ indicator GCaMP6f, we repeated widefield experiments in Pdgfrα^CreERT(+/-)^;Gcamp6f^Lox/Lox^ mice at 7 dpi and 14 dpi and obtained similar results (Supplementary Fig. 1). Finally, we confirmed that Ca²⁺ signaling was low in the healthy corpus callosum but strongly increased in demyelinating lesions (Supplementary Fig. 2), extending previous observations in myelin sheaths to all oligodendroglia^7^.

The dynamic behavior of oligodendroglial Ca^2+^ signaling during demyelination and remyelination has not yet been characterized in vivo. While our imaging experiments in acute slices revealed that these cells exhibit intrinsic Ca^2+^ activity in lesions, this approach does not enable monitoring their temporal evolution in the same animal. To address this limitation, we used longitudinal *in vivo* microendoscopy in Pdgfrα^CreERT(+/-)^;Gcamp6f^Lox/Lox^ mice during demyelination and remyelination while they explore an open field (Fig. 2a,b). Since the LPC model was incompatible with the multi-step procedure of microendoscopy, we used the cuprizone (CPZ) model, which induces progressive demyelination over several weeks (Fig. 2a)^6,13^. We consistently detected spontaneous, robust Ca^2+^ activity in oligodendroglial cells throughout the two last weeks of demyelination and across six weeks of remyelination (Fig. 2c-d; Supplementary Video 3). These findings provide direct in vivo evidence that oligodendroglial Ca^2+^ signals are sustained throughout both the demyelination and remyelination phases.

**Figure 2.**
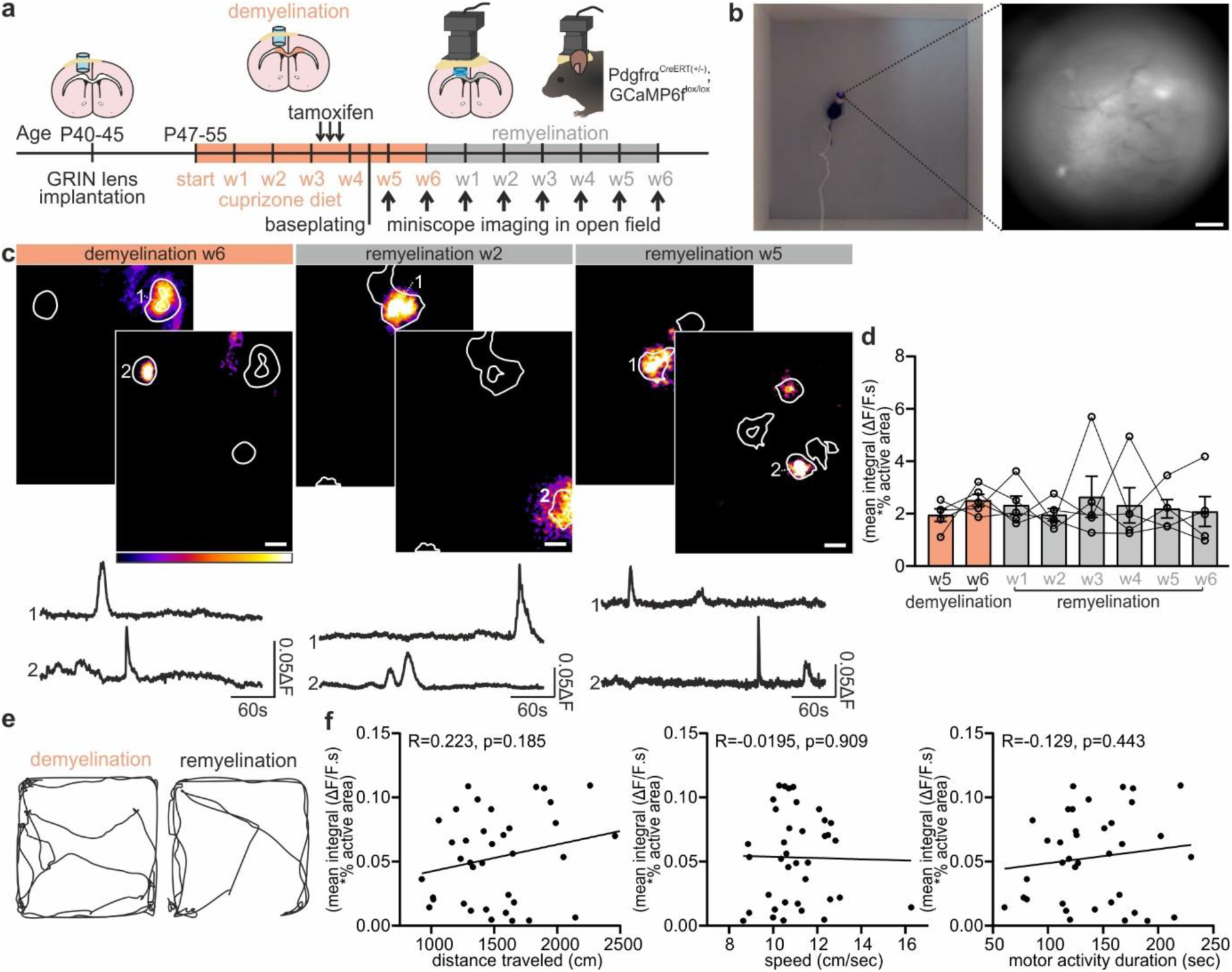
Longitudinal *in vivo* microendoscopy shows spontaneous, intrinsic Ca^2+^ activity during demyelination and remyelination. (**a**) Experimental timeline. Mice expressing GCaMP6f in oligodendroglia were implanted with a gradient index (GRIN) lens above the corpus callosum of the M1 motor cortex, ∼1.8 mm below the brain surface, and exposed to the CPZ diet for 6 weeks. UCLA Miniscope V4 baseplating was performed in the 5th week of CPZ demyelination and microendoscopy imaging was performed in weeks 5 and 6 of demyelination (orange) as well as during weeks 1 to 6 of remyelination (gray). (**b**) Miniscope recordings were performed in an open field and locomotion behavior was monitored. (**c,d**) Representative projection images of output by Occam with designated active ROIs (white lines) detected in several substacks as obtained with the Miniscope configuration of Occam (File S1)^6^. Two substack projections obtain from image stacks are shown per week (**c**). Ca^2+^ activity traces (**c**, bottom) and quantification (**d**) of *in vivo* oligodendroglial Ca^2+^ imaging during de-and re-myelination (not significant; one-way repeated measures ANOVA). (**e**) Open field locomotion maps during *in vivo* Ca^2+^ imaging, recorded under demyelination and remyelination conditions in the same mouse. (**f**) Correlations between Ca^2+^ signals and distance traveled, speed and motor activity duration (not significant Pearson correlations) during remyelination. Data represented as mean ± s.e.m. Scale bars 100 µm (**b**) and 20 µm (**c**).

Neuronal activity in the motor cortex changes dynamically during locomotion^14,15^. We thus explored whether Ca^2+^ signaling in oligodendroglia correlates with such neuronal activity in vivo. To this end, we recorded their Ca^2+^ signals in the corpus callosum directly underneath the M1 motor cortex while monitoring the locomotor behavior of mice and used locomotion as a proxy of neuronal network activity. Interestingly, oligodendroglial Ca^2+^ activity did not correlate with measures such as distance traveled, running speed, or duration of motor activity in the open field (Fig. 2e-f). These results align with our findings from acute slices and LPC-induced lesions which supports the conclusion that oligodendroglial Ca^2+^ signals in demyelinated lesions arise from intrinsic cellular mechanisms rather than follow the global neuronal network activity.

Altogether, our results demonstrate that oligodendroglial cells exhibit spontaneous, intrinsic Ca^2+^ activity appearing in the adult, within demyelinated lesions, and persisting throughout both demyelination and remyelination. We show that this signaling is largely independent of neuronal activity and network states associated with locomotion. These findings establish an intrinsic Ca²⁺ signaling behaviour in OPCs and OLs that is conserved across two distinct lesion models.

### Chemogenetic enhancement of oligodendroglial Ca^2+^ signals promotes oligodendrocyte regeneration

Our findings establish intrinsic Ca^2+^ signaling as a key regulator of the oligodendroglial response to injury. We next tested whether enhancing intrinsic Ca^2+^ signaling in these cells is sufficient to promote their regeneration. To address this question, we employed a chemogenetic strategy targeting oligodendroglia to selectively enhance their intrinsic Ca²⁺ signaling and investigate its causal effect on OPC and OL densities (Fig. 3a). Following LPC-induced demyelination of the corpus callosum, we directly assessed whether clozapine-N-oxide (CNO)-mediated activation of hM3D(Gq) receptor in acute brain slices of Pdgfrα^CreERT(+/-)^;Gcamp6f^Lox/Lox^;Hm3d(Gq)-mCherry^Lox/lox^ mice could potentiate Ca²⁺ signaling in oligodendroglia. We found that CNO exposure elicited sustained increases in Ca²⁺ signals within the lesion site in acute slices at 7 dpi, significantly potentiating the spontaneous Ca²⁺ activity (Fig. 3b). This establishes that our strategy was effective in selectively increasing intrinsic Ca²⁺ signaling in oligodendroglia after demyelination.

**Figure 3.**
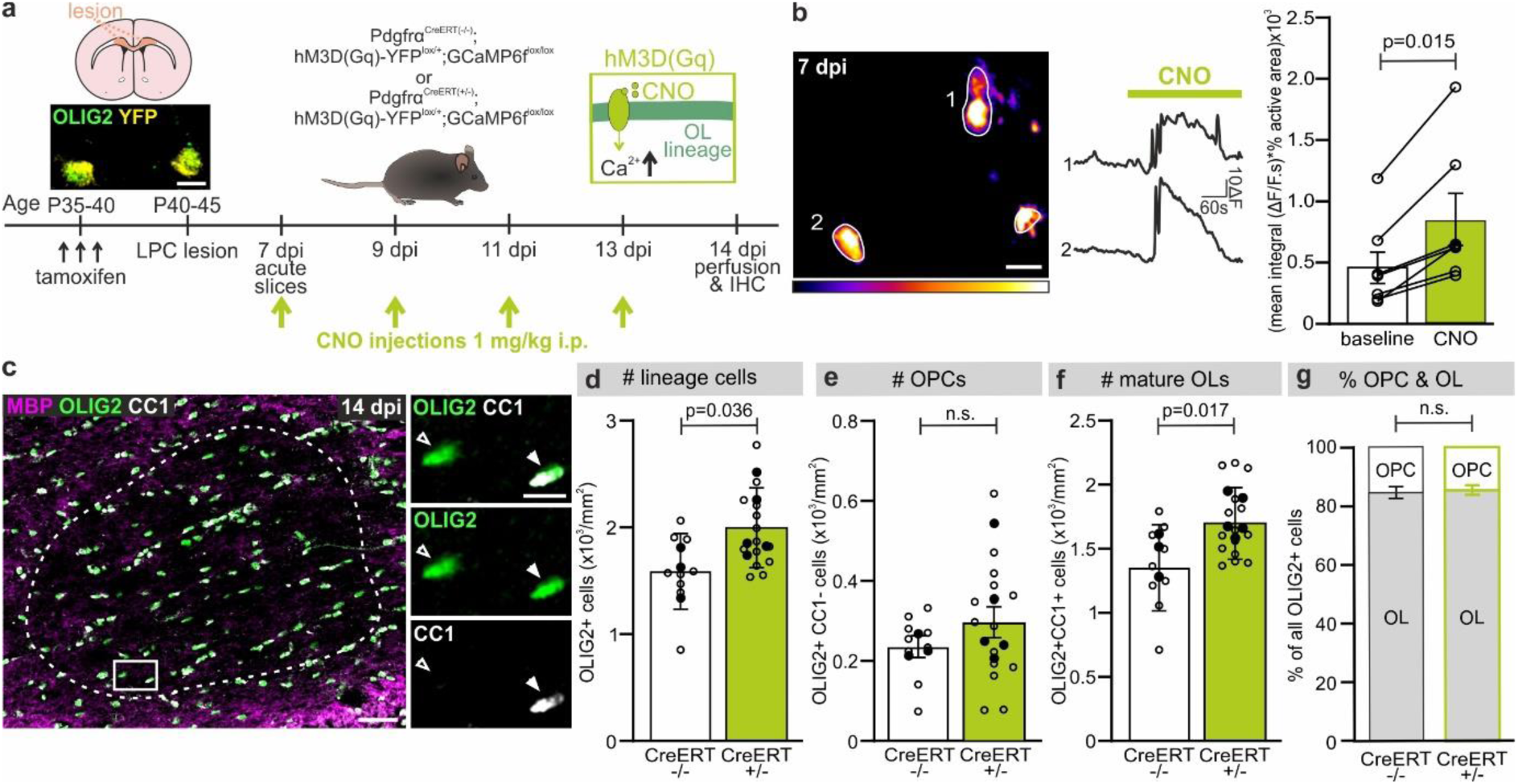
Enhancing intracellular Ca^2+^ signaling in oligodendroglial cells promotes lineage expansion in LPC-induced lesions. (**a**) Experimental timeline. Callosal LPC-induced lesions were induced in control mice (Pdgfrɑ^CreER(-/-)^;hM3D(Gq)-YFP^lox/+^;GCaMP6f^lox/lox^) and experimental mice (Pdgfrɑ^CreER(+/-)^;hM3D(Gq)-YFP^lox/+^;GCaMP6f^lox/lox^ mice) which express both GCaMP6f and the hM3D(Gq) receptor in oligodendroglial cells (Inset: OLIG2^+^YFP^+^ cells). Mice were either used to prepare acute slices at 7 dpi or received i.p. CNO injections (1 mg/kg) at 7, 9, 11 and 13 dpi before perfusion at 14 dpi. (**b**) Representative projection image output by Occam with designated active ROIs (white lines) and Ca^2+^ traces (middle) from a lesion at 7 dpi. CNO bath application significantly increased Ca^2+^ activity compared to baseline. Dot plots (right) showing significant Ca^2+^ activity increases in the presence of CNO in lesions (p-value from Wilcoxon matched-pairs signed rank test). (**c**) Representative image of immunofluorescence staining for MBP (magenta), OLIG2 (green) and CC1 (white) in a callosal lesion (dashed line). Insets from white rectangle shows an OLIG2^+^CC1^-^ OPC (open arrow) and mature OLIG2^+^CC1^+^ OL (closed arrow). (**d-f**) Densities of OLIG2^+^ lineage cells (**d**), OLIG2^+^CC1^-^ OPCs (**e**) and OLIG2^+^CC1^+^ OLs (**f**) in Cre^-/-^ (control) and Cre^+/-^ mice following CNO injections. (**g**) Percentages of OLIG2^+^ cells that are OPCs (OLIG2^+^CC1^-^) or mature OLs (OLIG2^+^CC1^+^) did not differ between CNO-treated mice with and without oligodendroglial hM3D(Gq) receptor expression. p-values from mixed linear models with type II likelihood ratio tests (**d-g**). Data is presented as mean ± s.e.m. with open circles representing slices and closed circles averages per mouse (**d,f**). Not significant n.s. Scale bars are 5 µm in (**a**), 10 µm in (**b**) and 50 µm and 10 µm in (**c**).

To evaluate the functional effect of enhanced Ca²⁺ signaling on OL regeneration, we induced LPC-induced lesions in mice and administered the brain-permeable CNO every other day between 7 and 14 dpi, a period during which OPCs start differentiating into OLs. Immunohistochemical analysis at 14 dpi revealed that *in vivo* chemogenetic activation of oligodendroglia significantly increased the number of OLIG2^+^ OL lineage cells (Fig. 3c,d) and OLIG2^+^CC1^+^ OLs in mice expressing the hM3D(Gq) receptor (Fig. 3f). The number of OLIG2^+^CC1^-^ OPCs was not significantly different (Fig. 3e), consistent with the small OPC pool typically observed at 14 dpi^16^, but the proportion of OPCs and OLs remained stable, indicating that differentiation rates were unchanged (Fig. 3g).

We conclude that potentiation of cell-autonomous Ca^2+^ activity promotes the expansion of the oligodendroglial lineage, resulting in increased numbers of mature OLs, without altering the efficiency of differentiation.

### Mitochondrial Ca^2+^ flux sustains oligodendroglia Ca^2+^ signals

We found that intrinsic Ca^2+^ signals promote OL regeneration, but the subcellular mechanisms that generate and sustain these signals during remyelination remain unknown. In astrocytes, intracellular Ca^2+^ dynamics depend on mitochondrial efflux^17^. Mitochondria have also been implicated in myelin sheath Ca^2+^ activity during development^7^, yet their potential role in coordinating the Ca²⁺ signals that govern OL regeneration is unknown. To address this question, we tested whether mitochondrial Ca^2+^ efflux sustains oligodendroglial activity by blocking the mitochondrial permeability transition pore (mPTP) with cyclosporin A in acute slices, in combination with rotenone to inhibit mitochondrial complex I, thereby enhancing the blockade of mPTP opening^7,17^. Strikingly, mPTP blockade abolished almost all oligodendroglial Ca^2+^ signals, identifying mitochondria Ca^2+^ efflux as the principal source of intrinsic intracellular Ca^2+^ activity in lesions (Fig. 4a).

**Figure 4.**
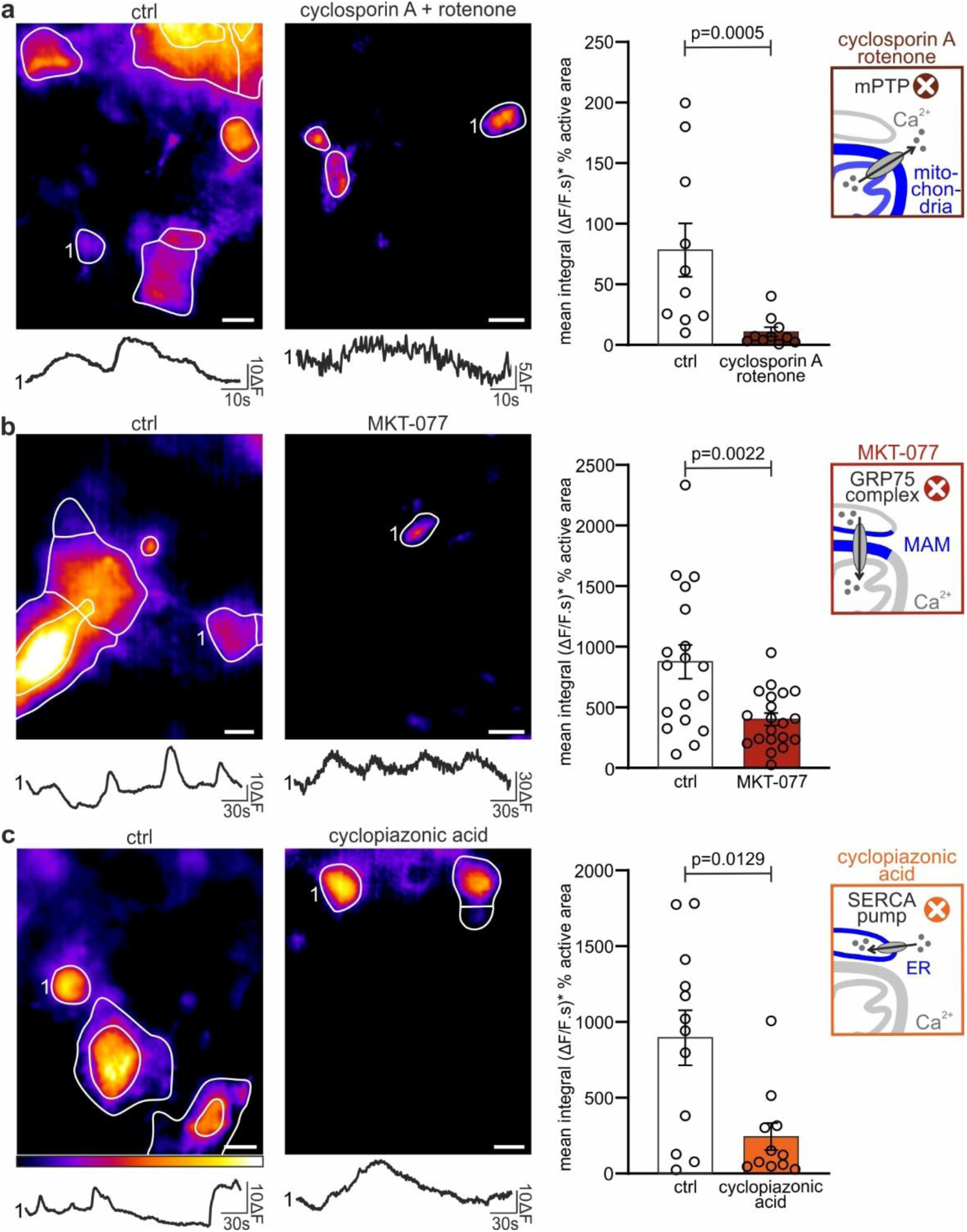
ER-mitochondrial Ca^2+^ flux sustains intrinsic oligodendroglial Ca^2+^ activity LPC-induced lesions. (**a,b,c**) Representative projection image outputs by Occam with designated active ROIs (white lines; left, top) and corresponding Ca^2+^ traces (left, bottom) from lesions at 7 dpi in control and after incubation with drugs for at least 30 min prior to recordings (see Online Methods). Dot plots of Ca^2+^ activity (right) under control conditions (white) and after pharmacological inhibition (color) of ER-mitochondrial pathway with 20 µM cyclosporin A and 10 µM rotenone (**a**), 10 µM MKT-077 (**b**) and 100 µM cyclopiazonic acid (**c**). p-values from two-tailed Mann-Whitney U tests (**a,c**) and two-tailed unpaired Student’s t test (**b**). Dot plots represent mean ± s.e.m. Scale bars 10 µm.

We next asked how the mitochondria Ca^2+^ store is replenished. One possibility is the Ca^2+^ transfer from the endoplasmic reticulum (ER) through the mitochondria-associated membrane (MAM). To test this, we incubated the slices with MKT-077, the pharmacological inhibitor of the GRP75 complex, which disrupts the ER-mitochondria coupling. MAM inhibition significantly reduced spontaneous Ca^2+^ activity in oligodendroglia (Fig. 4b), indicating that ER-to-mitochondria Ca^2+^ flux partially sustains intrinsic Ca^2+^ signals. To confirm the involvement of the ER, we blocked the SERCA pump with cyclopiazonic acid, preventing ER Ca^2+^ store replenishing. Prolonged incubation with cyclopiazonic acid caused a significant decrease in spontaneous oligodendroglial Ca²⁺ signals (Fig. 4c), supporting the conclusion that ER stores, in concert with MAM-mediated transfer, are essential for maintaining mitochondrial-dependent intrinsic Ca^2+^ signaling. Importantly, neuronal synaptic transmission was unaffected by cyclopiazonic acid or MKT-077 (Supplementary Fig. 3), indicating that the observed effects are intrinsic to oligodendroglia.

To test whether other pathways known to be implicated in oligodendroglial physiology might also contribute to these Ca^2+^ signals, we applied antagonists of voltage-gated K^+^ and Ca^2+^ channels, α1 adrenergic receptors, and TrkB receptors^8,18–20^. None of these drugs significantly reduced intrinsic oligodendroglial Ca^2+^ activity (Supplementary Fig. 4), suggesting that, while these pathways may influence different aspects of oligodendroglia biology, they are not required for preserving the spontaneous oligodendroglial Ca^2+^ activity that appears in demyelinated lesions. Together, these results identify the mitochondrial Ca²⁺ efflux, supported by ER-derived Ca²⁺ flow through MAMs, as the central mechanism sustaining intrinsic oligodendroglial Ca²⁺ activity in lesions, independent of previously described neuronal input or neuromodulatory pathways.

### Mitochondrial Ca^2+^ flux is required for OL regeneration

We next investigated the functional consequences of the blockade of mitochondrial Ca^2+^ flux on OL regeneration *in vivo*. To this end, we induced LPC-induced lesions in the corpus callosum and implanted an intraventricular mini-pump releasing MKT-077 to inhibit MAM Ca^2+^ transfer during early remyelination from 7 to 10 dpi (Fig. 5a). Prior to treatment, we confirmed that this intraventricular infusion effectively delivered compounds to the corpus callosum by demonstrating the diffusion of the 3000 MW Dextran Texas red tracer (Fig. 5b). Immunohistochemical analysis at 10 dpi revealed that MKT-077 treatment led to a significant decrease in the densities of OLIG2^+^ OL lineage cells (Fig. 5c,d) and mature OLIG2^+^CC1^+^ OLs (Fig. 5f). The proportion of differentiated OLs relative to OPCs was also reduced (Fig. 5g), indicating a defect in oligodendroglial differentiation and the establishment of an OL/OPC imbalance. However, this imbalance was not accompanied by the expansion of the OPC pool as the density of OLIG2⁺CC1⁻ OPCs at 10 dpi remained unchanged (Fig. 5e). This suggests that, in MKT-077-treated lesions, OPCs fail to compensate for the loss of differentiated OLs, leading to a persistent imbalance in the OL/OPC ratio. Because *in vivo* analysis was performed after the treatment period (7–10 dpi), these experiments could not directly capture the initial potential effects of MKT-077 on OPC proliferation itself. To address this question in a simplified system where oligodendroglial cells can be directly targeted without confounding tissue-level effects, we turned to a cell culture model. We employed the CG4 OPC cell line to examine the direct impact of MKT-077 on OPC survival and proliferation (Fig. 6a). MKT-077 exposure for 3 days *in vitro* (DIV) induced minimal apoptosis (5.75±1.44% of cells expressed the cleaved caspase-3⁺ marker in 0.5 µM MKT-077; n=3) and OPCs retained normal morphology (Fig. 6b). However, it strongly reduced OPC proliferation in a dose-dependent manner (Fig. 6b). Altogether, these results indicate that disruption of ER–mitochondrial Ca²⁺ coupling impairs OL differentiation *in vivo* and, by directly reducing OPC proliferation as confirmed *in vitro*, contributes to defective oligodendroglia repopulation at the lesion site.

**Figure 5.**
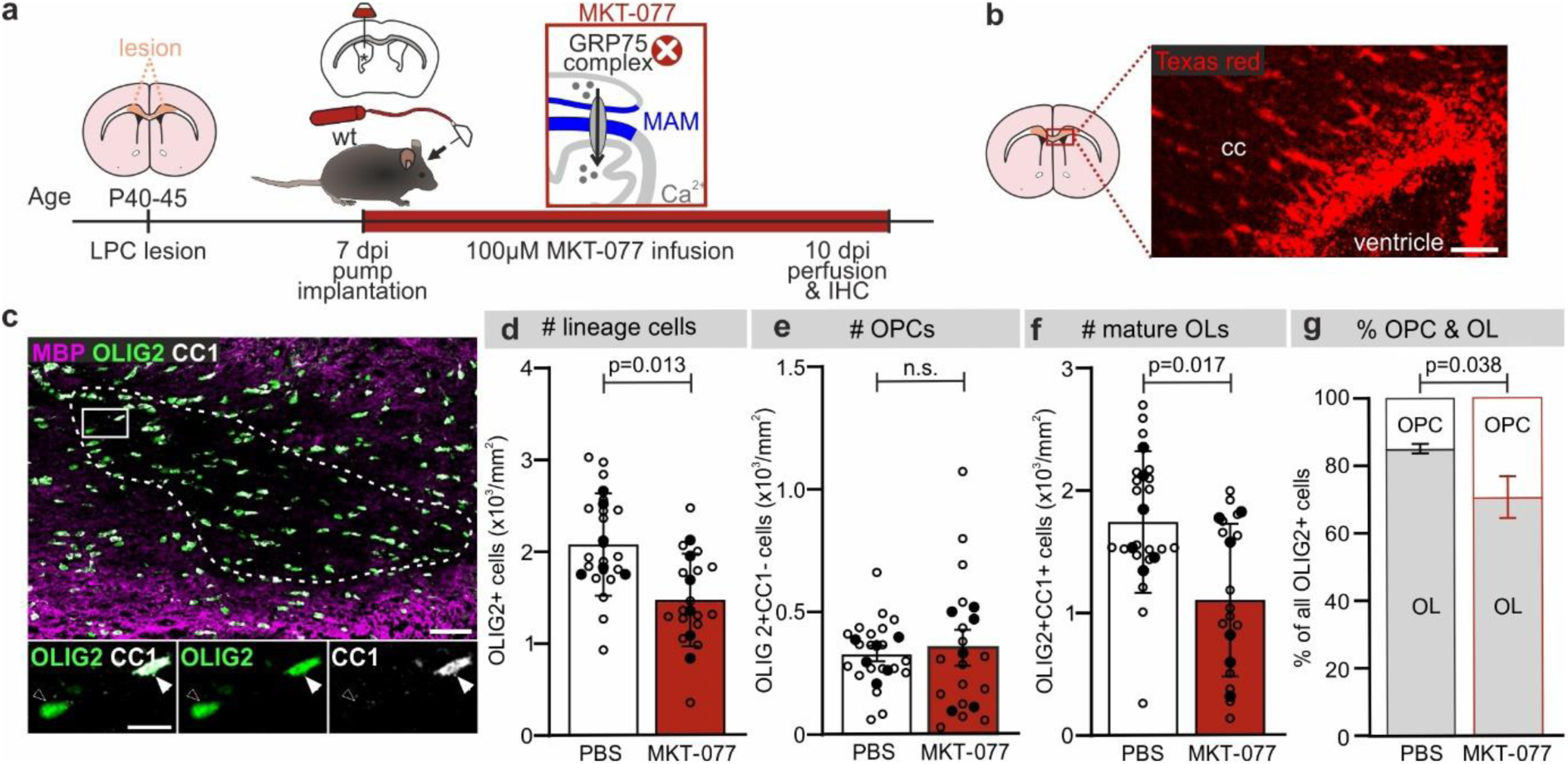
Inhibiting mitochondrial Ca^2+^ flux in oligodendroglial cells is detrimental to lesion repopulation and OPC differentiation. (a) Experimental timeline. LPC-induced lesions were induced in the corpus callosum of wild type mice and minipumps with PBS or MKT-077 delivered at a rate of 0.11 µL/hour were implanted intraventricularly at 7 dpi. Mice were infused from 7-10 dpi and perfused at 10 dpi. (b) Validation of experimental approach showing that 3000 MW Dextran Texas red dye infused in the ventricles *via* a minipump reaches the corpus callosum (cc). (**c**) Representative immunofluorescence image showing a lesion (dashed line) with MBP (magenta), OLIG2 (green) and CC1 (white). Insets (bottom) show OLIG2^+^CC1^-^ OPC (open arrow) and OLIG2^+^CC1^+^ mature OL (closed arrow). (**d-f**) Densities of OLIG2^+^ lineage cells (**d**), OLIG2^+^CC1^-^ OPCs (**e**) and OLIG2^+^CC1^+^ mature OLs (**f**) in lesions from control (white) and MKT-077-treated mice (dark red). (**g**) Percentages of OLIG2^+^ cells that are OPCs (OLIG2^+^CC1^-^) or mature OLs (OLIG2^+^CC1^+^) differ between mice in control and receiving MKT-077 infusions. P-values from mixed linear models with type II likelihood ratio tests. Data is presented as mean ± s.e.m with open circles representing slices and closed circles representing averages per mouse. Scale bars are 50 µm in (**b**) and 50 µm and 10 µm in (**c**).

**Figure 6.**
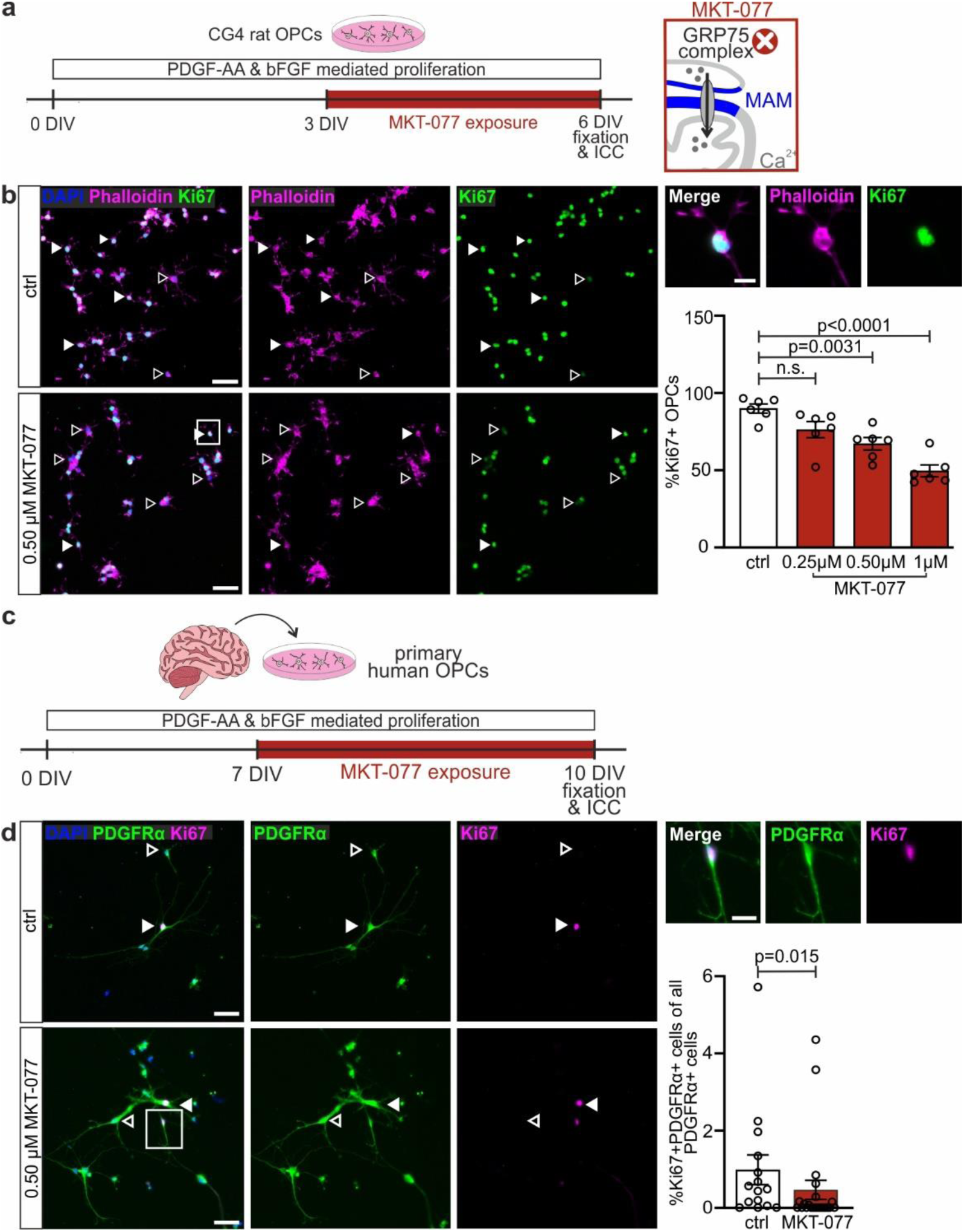
ER-mitochondrial contact sites for Ca^2+^ transfer are required for OPC proliferation. (a) Rat OPCs from the CG4 cell line were maintained under growth-factor stimulated proliferation condition through exposure to PDGF-AA and bFGF for 6 days *in vitro* (DIV). Cultures were exposed to either control conditions or 0.25 µM, 0.50 µM or 1 µM of MKT-077 for 3 DIV. (**b**) Representative images and quantification of cell nuclei (DAPI^+^; blue), actin filaments to expose cellular morphology (Phalloidin^+^; magenta) and proliferative cells (Ki67^+^; green). Non-proliferative OPCs (Ki67^-^; open arrows) and proliferative OPCs (Ki67+; closed arrows) are indicated. The highlighted region shows that proliferative OPCs under 0.50 µM MKT-077 exposure maintained normal morphology. Dot plots (right) of the % of Ki67+ OPCs are represented as mean ± s.e.m. P-values are from one-way ANOVA test followed by a Holm-Sidak’s multiple comparisons testing. Scale bars are 10 µm and 50 µm. (**c**) We obtained primary human OPCs *via* O4-mediated extraction from fresh tissue samples obtained during brain surgeries. Cells were maintained under growth-factor stimulated proliferation condition using PDGF-AA and bFGF for 10 DIV, and exposed to MKT-077 from 7 to 10 DIV. OPCs were fixed and stained at 10 DIV. (**d**) Representative images and quantification of cell nuclei (DAPI^+^; blue) OPCs (PDGFRɑ^+^; green) and proliferative OPCs (Ki67^+^; magenta). Non-proliferative OPCs (open arrows; PDGFRɑ^+^Ki67^-^) and proliferative OPCs (closed arrows; PDGFRɑ^+^Ki67^+^) are indicated. The highlighted region shows that proliferative OPCs under MKT-077 exposure maintained normal morphology. Data from 6 donors in the control (ctrl) group and 5 donors in the MKT-077 group. P-value from two-tailed Mann-Whitney U test. Dot plots (right) of the % of PDGFRɑ^+^Ki67^+^ OPCs are represented as mean ± s.e.m. (**d**) and n.s. not significant. Scale bars are 10 µm and 50 µm (**b,d**).

Since human oligodendroglia exist in a far wider spectrum of cell states than found in rodent oligodendroglia, we aimed to establish whether the mechanism described above in mitochondria is conserved across species. To determine the relevance of our findings for human OPCs, we employed primary human OPC cultures. We performed O4-mediated extraction of OPCs from freshly resected human brain tissue obtained from surgeries at Amsterdam UMC (see Online Methods and Supplementary table 1 for patient characteristics).

These human OPCs were maintained under proliferative growth factor exposure for 10 DIV, in the absence or presence of MKT-077 during 7 to 10 DIV, during which they retained normal morphology (Fig. 6c). Consistent with our observations in rodent CG4 OPCs, blocking the GRP75 protein at the MAM significantly reduced OPC proliferation (Fig. 6d), suggesting that the requirement for ER-mitochondrial contact sites in OL regenerative responses is conserved in humans.

In conclusion, these results demonstrate that mitochondrial-dependent Ca^2+^ signaling is an essential cell-intrinsic mechanism sustaining oligodendroglial activity that governs proliferation and differentiation during remyelination. Consistent with these observations, pharmacological inhibition of ER-mitochondrial contact sites for Ca^2+^ transfer in primary human OPCs similarly impaired OPC proliferation, suggesting that the underlying mitochondrial regulation of oligodendroglial lineage progression is conserved across species.

## Discussion

Our study uncovers a previously unknown role for cell-autonomous Ca²⁺ signaling in OL regeneration following demyelination. We demonstrate that spontaneous intrinsic Ca²⁺ activity appear in OPCs and OLs in demyelinated lesions and persist throughout both demyelination and remyelination, independently from neuronal network activity. Mechanistically, we identify mitochondrial Ca²⁺ efflux as the primary source of oligodendroglial Ca^2+^ signals. We further show that the mitochondrial Ca²⁺ store is replenished *via* ER-mitochondria contact sites, establishing an intracellular Ca²⁺ flux that is critical for OL regeneration. Disruption of this pathway impairs OPC proliferation and differentiation, revealing a function for mitochondrial Ca^2+^ flux in the regeneration of the OL lineage. Importantly, inhibition of ER-mitochondrial Ca^2+^ coupling similarly reduced proliferation in primary human OPCs, suggesting that mitochondrial regulation of oligodendroglial dynamics is a conserved feature across species and may represent a potential therapeutic target to treat demyelination-related disorders such as multiple sclerosis (MS).

In the zebrafish spinal cord, during CNS development, OPCs segregate into two functionally distinct subtypes showing different Ca²⁺ activity levels: one subtype is characterized by high Ca²⁺ signaling and greater proliferative capacity, while the other subtype is distinguished by lower Ca²⁺ activity but enhanced differentiation potential.^21^ Other *in vivo* studies have shown that Ca²⁺ dynamics in OLs regulate myelin sheath formation, stabilization, and remodeling, with localized transients predicting both sheath growth and retraction^11,22–24^. In rodents, Battefeld et al.^7^ reported that Ca^2+^ signaling in myelin sheaths is prominent during development but disappears with age. Together, these studies support a central role for Ca²⁺ signaling in lineage progression and myelin plasticity during development.

Interestingly, several studies have shown that adult OPCs can undergo, to some extent, molecular and functional rejuvenation in response to injury^3,25^, including reactivation of Ca^2+^ transients in myelin sheaths during remyelination^7^. We demonstrate that these reactivated Ca²⁺ signals in the entire OL lineage are not merely a marker of injury, but play an essential functional role in OL regeneration. These results support the view that OL lineage cells can regain a developmentally permissive state during repair, with intracellular Ca²⁺ dynamics acting as a key actor governing their regenerative potential. Although our two-photon microscopy data failed to single out any Ca²⁺ signaling profile corresponding to either OPC or OL subpopulations, the potential for the existence of such functionally distinct subpopulations in lesions needs further investigation.

Myelination is now recognized as both an activity-dependent and activity-independent neuronal process. While neuronal activity can accelerate and enhance myelination, it is not strictly required as myelination can also occur without it^26^. In recent years, several neuronal pathways have been implicated in modulating oligodendroglial Ca²⁺ dynamics and lineage behavior, including K^+^-dependent signaling *via* Kir4.1 channels^18^, noradrenergic α-adrenoceptor activation^8,9^, and BDNF-TrkB signaling^20^. However, our findings reveal that the Ca^2+^ signals governing oligodendroglial regeneration after demyelination persist independently of these pathways. Blocking neuronal firing, K^+^ or Ca^2+^ channels, α₁-adrenoceptors, or TrkB receptors had no effect on spontaneous Ca^2+^ signals in demyelinated lesions, indicating that oligodendroglial signaling is maintained through an intrinsic, neuronal activity-independent mechanism. These results position mitochondrial Ca²⁺ efflux as a key component of OL regenerative signaling that regulate both proliferation and differentiation.

Studies in OPCs and OLs highlight a remarkable diversity of Ca^2+^-dependent mechanisms during development or after demyelination. *In vivo* imaging in zebrafish has shown that the frequency and amplitude of Ca^2+^ transients in OLs have distinct consequences: high-frequency events promote sheath elongation, whereas long-duration, high-amplitude transients trigger its retraction^11,23^. Complementary studies in mice have revealed that local Ca²⁺ transients in nascent myelin sheaths regulate cytoskeletal organization without directly affecting OL differentiation^27^. Genetic manipulations further illustrate this complexity: the deletion of Cav1.2 channels impairs remyelination by reducing OPC proliferation and OL generation^19^, yet in healthy adults, it causes only a transient, region-specific OPC loss without impacting overall OL production^28^. Our results add to this mechanistic diversity by identifying ER–mitochondria Ca²⁺ flux as the basis of intrinsic Ca²⁺ activity and OL regeneration in demyelinated lesions. Mitochondrial Ca²⁺ signaling thus becomes part of a highly adaptable signaling repertoire that enables oligodendroglia to adapt their proliferation-and differentiation-promoting responses to regenerative demands.

Mitochondria-based processes are known to regulate neural stem cell survival, proliferation, and differentiation in both embryonic and adult neurogenesis, often through (but not limited to) metabolic plasticity^29–31^. In myelinating cells, metabolic plasticity is achieved by mitochondrial Ca^2+^ flux, which modulates the mitochondrial membrane potential and facilitates ATP production^32^. Moreover, in OPCs mitochondrial networks are highly dynamic and mostly located in their processes, while during differentiation they expand and relocate to the soma^33^. These mitochondrial changes during OPC differentiation are associated with metabolic modifications which are sensitive to cellular stress^34,35^, highlighting their potential role during regeneration. We show that ER-mitochondrial contact sites mediating Ca^2+^ transfer is required to initiate proliferative processes in both rodent and human OPCs. Further research is necessary to establish whether the differentiation process depends on metabolic changes induced by mitochondrial Ca^2+^ flux. Our results highlight a signaling role for mitochondria, revealing its key function beyond their bioenergetic role. While our data point to a central role of Ca^2+^ signaling in OL regeneration, a contribution from mitochondrial metabolic pathways remains possible and needs further investigation.

Human oligodendroglia exist in a far wider spectrum of cellular and maturation states than found in rodent oligodendroglia, particularly in the context of demyelinating diseases such as MS^36^. Therefore, it is important to establish whether findings from rodent OL regeneration studies are relevant to the human condition. Our findings show that, even if OL may exist in a wider spectrum of cellular and maturation states in humans, the intrinsic processes that drive proliferation seem to be conserved across species. We here demonstrate that functionally, mitochondrial Ca^2+^ flux underpins the initiation of OPC proliferation in human OPCs as well as in rodent OPCs. Mitochondrial dysfunction is increasingly recognized as a key factor in the remyelination failure during disease progression in MS. In patient tissue and animal models, oligodendroglial mitochondria show abnormal morphologies, an impaired respiratory chain function, and disrupted Ca^2+^ homeostasis, all of which arrest OPC maturation and limit repair^37–39^. The effects of these alterations, ranging from respiratory deficits, increased oxidative stress, disrupted mitochondrial dynamics, and altered mitochondrial DNA integrity, contribute to cellular energy failure and neurodegeneration. By examining demyelinated lesions, we demonstrate that, in oligodendroglia, mitochondrial Ca^2+^ efflux, followed by Ca^2+^ store replenishing *via* ER–mitochondrial contact sites, sustains the intrinsic Ca^2+^ signaling required for OPC proliferation and differentiation during early remyelination. Our findings suggest that when mitochondrial Ca^2+^ flux is compromised, the regenerative program cannot be maintained, contributing to the differentiation block characteristic of MS. Thus, our work points to mitochondrial Ca^2+^ signaling as a promising therapeutic target in myelin-related disorders such as MS.

## Online Methods

### Transgenic mice

All experimental procedures adhered to the European Union and institutional guidelines for the care and use of laboratory animals. Ethical approval was obtained from both the French Ethical Committee for Animal Care of the University Paris Cité (Paris, France) and the Ministry of National Education and Research (Authorizations N°13093-2017081713462292 and 47940-2023101817084097).

Experiments were conducted using male and female transgenic mice, including Pdgfrα^CreERT(+/-)^;Gcamp6f^Lox/Lox^, Pdgfrα^CreERT(+/-);^Gcamp5-tdTomato^Lox/Lox^ transgenic adult mice obtained by crossing as described in Maas et al.^6^, and NG2^CreERT2(+/-)^;Gcamp3f^Lox/Lox^ obtained by crossing as in Balia et al.^40^. Pdgfrα^CreERT(+/-)^;Gcamp6f^Lox/Lox^;hM3D(Gq)-YFP^Lox/+^ were obtained by crossing Pdgfrα^CreERT(+/-)^;Gcamp6f^Lox/Lox^ mice and B6N;129-Tg(CAG-CHRM3*,-mCitrine)1Ute/J mice (The Jackson Laboratory, RRIID:IMSR_JAX:026220). Wild type C57BL6J were obtained from Janvier and Charles River laboratories. Genotyping was performed by PCR using primers for Cre or NG2-Cre^41^. All animals were housed under standard laboratory conditions, including ad libitum access to food and water, a 12-hr light/dark cycle, a controlled average temperature of 21°C and a relative humidity of 45%.

### Intracranial LPC injections

Intracranial injections of LPC (Sigma-Aldrich, Cat#62962) were done to induce a focal demyelination of corpus callosum in all mouse strains, as previously described^6^. Mice were deeply anesthetized with ketamine/xylazine (0.1/0.01 mg/kg, i.p.) and fixed in a stereotaxic frame (Kopf Instruments, Cat#940). Once positioned, they received a subcutaneous injection of buprenorphine (0.1 mg/kg) for analgesia. Throughout the surgical procedure, mice were maintained on a 37°C heating blanket to regulate body temperature, and ophthalmic dexpanthenol gel (Ocry-gel, TVM Dômes Pharma, France) was applied to the eyes to prevent dehydration. The skin of the head was disinfected with betadine, and lidocaine was applied locally before making the incision to expose the skull. Two small holes were drilled at the following coordinates: +1.40 mm anterior from bregma and ±0.95 mm lateral from the midline. Bilateral injections of 1.6 μL of LPC (10 mg/mL in filtered PBS) were administered at a depth of-1.70 mm. The skin of the skull was sutured to close the wound.

### Cuprizone treatment and *in vivo* microendoscopy

One week prior to the start of cuprizone (Sigma Aldrich, CAS: 370-81-0) treatment, *Pdgfrα^CreERT(+/-)^;Gcamp6f^Lox/Lox^* mice (P35-P42) were subjected to GRIN lens implantation surgery as previously described^6^. Briefly, a gradient index (GRIN) lens (1 mm diameter, 4 mm length; Inscopix, Cat#1050-004595) was implanted at a depth of 1.60 mm at coordinates +1.40 mm anterior to bregma and +1 mm lateral to the midline. At P42-P47, animals were given a diet consisting of 40% of cherry-flavored nutragel (Plexx, Cat#4798-KIT) and 60% MilliQ-water for 4-7 days. Over the next six weeks, the nutragel was supplemented with 0.3% CPZ to induce demyelination. The diet was refreshed daily, except on weekends, and the mice were weighed 2 to 3 times a week. The food was prepared every 4 days and kept at 4°C. In the fourth week of CPZ treatment, i.p. injections of tamoxifen were administered to induce GCaMP6f expression in OL lineage cells. During the fifth week of CPZ treatment, animals were subjected to a baseplating procedure as previously described^6^. Briefly, a baseplate was cemented onto the skull over the GRIN lens and positioned to allow clear imaging of the brain surface with the UCLA miniscope v4 (Open Ephys, Cat#OEPS-7407). *In vivo* Ca^2+^ imaging was performed on freely moving mice placed in a 50 cm × 50 cm x 45 cm open field arena (50 lux illumination) using the miniscope v4 at a frame rate of 10 fps for 10 min, while locomotor activity was synchronously monitored. Ca^2+^ signals were recorded from the fifth week of CPZ treatment to the sixth week of remyelination.

To monitor locomotor activity during *in vivo* Ca^2+^ recordings, mouse movement was recorded on freely moving mice with an infrared camera and analyzed with the ViewPoint behavioral analysis software (version 5.31.0.8, ViewPoint, Behavior Technology, France) between the 5th week of CPZ treatment and the 6th week of remyelination. Quantitative data including distance travelled, speed and motor activity duration were calculated by the software.

### Tamoxifen and CNO injections

Adult *Pdgfrα^CreERT(^*^±);^*Gcamp5-tdTomato^Lox/Lox^ and Pdgfrα^CreERT(^*^±)^*;Gcamp6f^Lox/Lox^*mice were intraperitoneally injected with tamoxifen (MP Biomedicals Germany GmbH, CAS: 10540-29-1) to induce the expression of GCaMP5 or GCaMP6f in OPCs and their progeny, respectively. For *Pdgfrα^CreERT(^*^±)^*;Gcamp6f^Lox/Lox^;hM3D(Gq)^Lox/+^* mice, tamoxifen injections were administered to express GCaMP6f and hM3D(Gq) receptors in OPCs and their progeny. In all cases, 1 mg tamoxifen was administered once per day for three consecutive days (1 mg in 100 µL of miglyol 812; Caesar & Lorentz GmbH). For mice with LPC-induced demyelination, tamoxifen was administered from 6 to 4 days prior to LPC injection (from P35-40). For mice undergoing CPZ treatment, tamoxifen injections began during the fourth week of the CPZ treatment (P60-P67). To induce GCaMP3 expression in *NG2^CreERT2(^*^±)^*;Gcamp3^Lox/Lox^* mice, 4-OHT (Hello BioTocris Bioscience) was administered to pups from P3 to P5 which ensured a high level of recombination in the adult without targeting neurons^41^. To activate hM3D(Gq) receptors *in vivo*, CNO (HelloBio, Cat#HB6149) was administered intraperitoneally. *Pdgfrα^CreERT(+/-)^;Gcamp6f^Lox/Lox^;hM3D(Gq)^Lox/+^* and *Pdgfrα^CreERT(^*^-/-)^*;Gcamp6f^Lox/Lox^;hM3D(Gq)^Lox/+^*mice received CNO at a dose of 1 mg/kg, a relatively low dose compared to studies on other glial cells^42^.

### *Ex vivo* widefield calcium imaging

Acute coronal slices (300 μm) were prepared as previously described^16^. Demyelinated lesions were visualized using DIC microscopy as brighter areas in the corpus callosum. The *ex vivo* widefield Ca^2+^ imaging of GCaMP5-and GCaMP6f-expressing cells was performed as previously described^6^ with an Olympus BX51 microscope equipped with a 40X fluorescent water-immersion objective. Imaging was performed at RT using an extracellular solution containing (in mM): 126 NaCl, 2.5 KCl, 1.25 NaH2PO4, 26 NaHCO3, 20 glucose, 5 pyruvate, 3 CaCl2 and 1 MgCl2 (95% O2, 5% CO2) and conducted in the corpus callosum with demyelinated lesions at either 7 dpi or 14 dpi. Several pharmacological and chemical agents were bath applied before and during recordings. Pre-recording bath applications included the following: 1 µM TTX, 200 µM Nickel and 200 µM Cadmium, 100 µM Barium for 5 minutes, 100 µM cyclopiazonic acid for 30 min, 10 mM TEA and 4 mM AP4 for 10 min and the cocktail of 100 µM AP5, 10 µM NBQX, 10 µM SR-95531, 5 µM CGP55835 and 20 µM MTEP for 20 minutes.

In some experiments, slices were pre-incubated with chemical agents for longer durations and in 3-5 mL solution: 20 µM cyclosporin A and 10 µM rotenone for at least 45 minutes, 1 µM cyclotraxin B for 2 hours, 10 µM MKT-077 (MedChem Express, Cat#HY-15096) for at least 45 minutes. For Ca^2+^ imaging in slices from *Pdgfrα^CreERT(^*^±)^*;Gcamp6f^Lox/Lox^;hM3Dq^Lox/+^*mice, 10 µM CNO was bath applied 2 min after starting the recording.

### *Ex vivo* two-photon Ca^2+^ imaging

Two-photon Ca^2+^ imaging of GCaMP5^+^ OPCs and OLs was performed in acute slices of demyelinated lesions, as previously described^6^. Briefly, imaging of individual cells was conducted on acute slices containing LPC-induced demyelinated lesions in the corpus callosum, using a two-photon laser scanning microscope operating in frame mode (150–250 ms per frame). Recordings were made for 99 s with custom-made software (Lab-VIEW, National Instruments, RRID:SCR_014468) and analyzed with Occam and post-prOccam programs as previously described^6^. The identification of OPCs and OLs was based on morphological criteria: OPCs were characterized by smaller, round somata and thin processes, while OLs had a larger soma and principal processes often aligned with axons. Only cells whose processes were unambiguously connected to the soma were included in the analysis.

### *Ex vivo* and *in vivo* Ca^2+^ imaging analysis

Ca^2+^ recordings were analyzed using the Occam and post-prOccam software programs (GNU GPLv3+ license; code repository at: https://gitlab.com/d5674/occam), that perform the automatic analysis of Ca^2+^ signals in OL lineage cells, as detailed^6^. In this report we used the software capabilities in the three widefield, two-photon and miniscope configurations. Quantitative data, including signal intensity and percentage of active area, were calculated by post-prOccam. For analysis of *in vivo* Ca^2+^ signaling, the last 5 consecutive minutes out of 10 recorded minutes were selected for the analysis, unless the baseline of Ca^2+^ signals was not constant. When the baseline was unstable, we chose 5 consecutive minutes with a constant baseline. For the longitudinal *in vivo* analysis described in this report we implemented in the Occam software a new filter for background homogenization that is specific for the miniscope configuration where it enables the correction for occasional light changes due to lens diffraction. This new filter has now been integrated into the open source software Occam package (v 1.0.6; see the new configuration file for the minsicope configuration in File S1).

### Patch-clamp recordings in acute slices

For patch-clamp recordings in the motor cortex, eEPSCs and eIPSCs of pyramidal neurons in control, in the presence of 100 µM cyclopiazonic acid (pre-incubation for at least 30 min) or 10 µM MKT-077 (pre-incubation for at least 45 min) were respectively evoked at −70 mV and 0 mV by a monopolar electrode (glass pipette) placed in layer V (100 ms pulse, 10 V, 0.2 Hz; Iso-Stim 01D, npi electronic GmbH, Tamm, Germany), with a CsMeS-based intracellular solution containing (in mM): 125 CsCH3SO3H, 5.4-aminopyridine, 10 tetraethylammonium chloride, 0.2 EGTA, 0.5 CaCl_2_, 2 MgCl_2_, 10 HEPES, 2 Na2-ATP, 0.2 Na-GTP and 10 Na_2_-phosphocreatine (pH ≈ 7.3). Recordings were made without series resistance (Rs) compensation. Rs was monitored during recordings and cells showing a change of more than 30% were not included in the analyses. Potentials were corrected for a junction potential of−10 mV. Acquisition was obtained using Multiclamp 700B and pClamp version 10.7 software, filtered at 4 kHz and digitized at 20 kHz. EPSCs and IPSCs were analyzed off-line using pClamp version 10.7 software (Molecular Devices, RRID:SCR_011323).

### *In vivo* MKT-077 infusions

*In vivo* MKT-077 infusions were performed in wild type C57BL6J mice. Minipumps (Alzet Cat#1004 with brain infusion kit) were filled with MKT-077 (100 µM) or sterile PBS and incubated in sterile PBS (pH 7.2) at 37°C for 3 days. The pump implantation surgery was performed in LPC lesioned mice at 7 dpi. A hole was stereotaxically drilled to implant a cannula into the right lateral ventricle using coordinates: −1.0 mediolateral, −0.50 anterior-posterior, −2.50 dorsoventral relative to bregma for local delivery. The cannula was then attached to the micro-osmotic pump containing MKT-077 or PBS via a catheter and placed subcutaneously between the shoulder blades. Infusion lasted for three days (7-10 dpi) at a flow rate of 0.11 µL per hour. Mice were perfused at 10 dpi.

### Tissue immunostaining and imaging

Mice were perfused intracardially with 2% paraformaldehyde (PFA). Brains were kept overnight in 2% PFA and then incubated in 30% sucrose in PBS or in a gradient of 10-20% sucrose at 4°C for around 3 days for cryoprotection. Brains were then embedded in OCT (Microm Microtech), frozen using isopentane and stored at-70°C. Coronal slices were cut with a cryostat (10 μm; Leica CM3050 S Cryostat), permeabilized in 4% bovine serum albumin (BSA, Sigma-Aldrich Cat#A7906) 0.1% triton X-100 and 10% normal goat serum (NGS, Sigma-Aldrich Cat#S26) for 1 hour and incubated with primary antibodies overnight. Immunostainings against OLIG2, CC1, and MBP were performed with rabbit anti-OLIG2 (1:400, RRID:AB_570666, Millipore), mouse anti-CC1 (1:100, RRID:AB_2057371, Millipore), and rat anti-MBP (1:100, RRID:AB_305869, Abcam). Secondary antibodies were Alexa-fluor 405 (1:200, ThermoFischer Scientific, RRID: AB_1307537), Alexa-fluor 680 (1:200, ThermoFischer Scientific, RRID: AB_2535723) and Alexa-fluor 546 (1:200, Thermo Fisher Scientific, RRID: AB_2534125) and were incubated for 2 hr at RT. Coverslips were washed with PBS 1X and mounted with Fluoromount-G (Southern Biotech, Cat# 0100-01). Confocal images were acquired with a ×63 oil objective of an SP8 confocal microscope (Leica TCS SP8 STED 3D) and the LAS X software (version 3.5.7.23225, Leica Microsystems, RRID: SCR_013673).

All cell counting was performed using the CellCounter plugin in Fiji/ImageJ2 (NIH, version 1.53i, RRID:SCR_002285)^43^ by experimenters that were blind to the experimental condition, ensuring unbiased analysis. The LPC-induced lesion was characterized as the region with a diminished MBP intensity in the corpus callosum. Cell densities were calculated as the number of cells per mm^2^ in the lesion area and the percentages of OPCs (OLIG2^+^CC1^-^) and OLs (OLIG2^+^CC1^+^) were defined as the number of cells divided by the total number of OLIG2^+^ lineage cells.

### CG4 OPC cell cultures

OPC rat cell line CG4^44^ was cultured in an N1 medium containing: DMEM/F12 (Gibco, Cat# 31331-028), 1 mg/mL insulin (Gibco, Cat# 12-585-014), 1x N1 mix (1 ng/mL selenite (Sigma-Aldrich, Cat# S5261), 3.22 mg/mL putrescine (Sigma-Aldrich, Cat# P5780) and 1 mg/mL transferrin (Sigma, Cat# T81558) in HBSS w/o Ca^2+^/Mg^2+^) and 124 ng/mL progesterone (ThermoFisher Scientific, Cat# 225650050). OPCs were maintained in proliferation medium, N1 medium supplemented with 25 ng/mL bFGF (Preprotech, Cat# AF-100-18B), 10 ng/mL PDGF-AA (Preprotech, Cat# 100-13A) and 0.01 µg/mL biotin (Sigma, Cat# B4639), for 6 DIV and from 3-6 DIV either DMSO or 0.25µM, 0.50µM or 1µM MKT-077 were applied.

### Primary human OPC cell cultures

Resected brain tissue from standard of care brain surgeries was processed following previous methods^45^. The tissue used in this study was ‘en route’ towards the resection target and therefore leftover. All patients gave written informed consent for the use of their data and leftover tissue before participation (reference 2010/126, approved 26 may 2010). Patient characteristics can be found in Supplementary table 1. Upon collection, tissues were kept at 4°C in Hibernate-A medium (ThermoFisher Scientific, Landsmeer, The Netherlands, Cat# A1247501). Within 30 minutes, samples were cut into pieces of < 1mm^3^ and digested with 0.25% trypsin (Gibco, Life Technologies, Paisley, UK, Cat# 11538876) for 30 min at 37°C under agitation (70 rpm) followed by an incubation with 20 U/mL DNase I (Roche Diagnostics GmbH, Mannheim, Germany, Cat# 4536282001) for 10 min. Fetal bovine serum (Thermo Fisher Scientific, Hampton, NH, USA, Cat# A31605) was added to the tissue homogenate and the homogenate was centrifuged for 10 min at 300g. Two washes were performed followed by a filtration through a 100 μm nylon cell strainer (10 min, 300g) with GNK-BSA buffer containing: 2.2 mg/mL D-Glucose monohydrate (Sigma-Aldrich, Cat# G8769-100ML), 3.3 mg/mL BSA (Roche, Cat# 10 735 086 001), dissolved in PBS 1X (Corning, New York, USA, Cat# CLS431751). Next, Percoll (GE Healthcare Bio-sciences AB, Uppsala, Sweden, Cat# GE17-0891–01) density gradient centrifugation was used to remove myelin and debris (30 min at 3200 x g at 4°C). The turbid phase containing the cellular fraction (the second fraction with the lowest density) was collected and washed in GKN-BSA buffer. The cell pellet was resuspended and washed in a sorting buffer (PBS 1X, 1% FBS, 2 mM EDTA (ThermoFisher Scientific, Cat# AM9260G)). Microglia cells were removed by magnetic cell separation (MACS) with α-CD11b microbeads (1:10, Miltenyi Biotec, Cat# 130-093-634) in sorting buffer for 30 min at 4°C and purified using a magnetic column. The negative fraction was collected, centrifuged 5 min 300 x g at 4°C and the pellet was incubated with α-O4 microbeads (Miltenyi Biotec, Cat# 130-094-543) for 30 min at 4°C and purified using a magnetic column to extract OPCs. Primary human OPCs were collected in OPC proliferation medium (DMEM/F12/GlutaMax medium (Gibco, Life Technologies, Paisley, Scotland, UK, Cat# 31331-028) supplemented with 1X B-27 with vitamin A (Gibco, Life Technologies, Cat# 17504044), 1X N-2 supplement (Gibco, Life Technologies, Cat# 17502048), 1 µg/mL laminin (Sigma Aldrich, Cat# L2020), 10 µg/mL bFGF, 25 µg/mL PDGF-AA (ReproTech, ThernoFischer Scientific, Cat# AF-100-18B and 100-13A) and 1% penicillin-streptomycin (Sigma-Aldrich, Cat# P4458)) before seeding. Isolated OPCs were seeded in 24-well plates pre-coated with poly-L-ornithine (100 µg/mL, Sigma-Aldrich, Cat# P3655) at a density of 15,000 cells per well. After 48 hours, the medium was refreshed. OPCs were kept for 10 DIV and exposed to DMSO or MKT-077 (0.25 µM or 0.50 µM) from 7-10 DIV.

### CG4 OPCs and primary human OPC immunostaining

OPCs were fixed with 2% PFA for 15 min at RT, then washed 3 times with PBS. Cells were permeabilised and blocked simultaneously with 5% normal donkey serum (Jackson ImmunoResearch, RRID: AB_2337258) and 0,1% Triton X-100 (Merck, Cat#1.08603) in PBS for 5 min, followed by 1.5-hour incubation with the primary antibodies in 5% donkey serum in PBS. After 3 washes of PBS 1X, secondary antibodies and DAPI (1µg/ml, Sigma-Aldrich, D9542) were incubated for 45 minutes. Immunocytochemistry of primary human OPCs included primary antibodies PDGFRɑ (R&D Systems, RRID: AB_354459, 1:100) for OPCs and Ki67 (Abcam, RRID: AB_443209, 1:400) for proliferating cells with secondary antibodies Alexa fluor 647+, 488 or 594 (Thermo Fisher, RRID: AB_2534102, 1:1000). Immunocytochemistry of CG4 OPCs included Ki67 (Abcam, Cat# ab15580, 1:400) for proliferating cells with secondary antibody Alexa fluor 488 (Thermo Fisher, RRID: AB_2534102, 1:1000) and Phalloidin (Invitrogen, Cat# R415, 1:500) to reveal morphology. Coverslips were washed with PBS 1X and mounted with Fluoromount-G (Southern Biotech, Cat# 0100-01). Images were acquired with a slide scanner SlideView VS200 (Olympus) with a 20X objective, 0.8NA, final resolution = 0.345 µm/pixel. Acquired images were processed and analysed with QuPath 5.0.1^46^.

Cell counting was conducted using the “cell detection” plug-in in Qpath, with DAPI serving as the reference fluorescence channel (threshold=1000, min size=10, max size=300). An individual classifier was trained for each channel by using the “train object classifier” plug-in, applying a RTress classifier (max trees=150, epsilon=0.01) and creating a composite classifier. For the CG4 OPCs, the percentage of Ki67^+^ cells over the total number DAPI^+^ cells was calculated. For the primary human OPCs, the proliferative capacity was assessed using the ratio of Ki67^+^PDGFRα^+^ cells over the total amount of PDGFRα^+^ cells.

## Statistical analyses

All data are represented as mean ± s.e.m. Statistical analyses were performed using GraphPad Prism (version 10; GraphPad Software Inc., USA), and JASP (version 0.16.4; JASP Team, 2022, Amsterdam, The Netherlands). Each group of data was first subjected to Shapiro-Wilk normality test. According to the data structure and variance equality, two-group comparisons were performed using either parametric or non-parametric tests. Specific statistical tests used in each comparison can be found in figure legends. Differences were considered significant when p< 0.05.

## Data Availability

The data generated and analyzed during this study are included in the manuscript and supplementary data. The raw datasets are available for research purposes from the corresponding author upon request.

## Code Availability

Occam and post-prOccam are open source analysis softwares and freely available (GNU GPLv3 + license; code repository at: https://gitlab.com/d5674/occam).

## Acknowledgements

We thank the NeurImag facility and the animal facility of IPNP and their funding sources (Fédération pour la Recherche Médicale, Fondation Leducq). NeurImag is a member of the France BioImaging Infrastucture (ANR-10-INSB-04). We also thank the imaging core facility of Amsterdam UMC. We would also thank Beata Turanska and Julie Cognet for their technical assistance, Serge Charpak and Yannick Goulam for their help with the two-photon microscope and Brahim Nait-Oumesmar for providing us with the CG4 cell line. Finally, we would also like to thank neurosurgeons Philip De Witt-Hamer and David Noske who helped us obtain human brain tissue for primary human OPC cultures and Marike van Lingen and Brigit Thomassen for handling logistics during this process.

## Author contributions

D.A.M. and R.B-V. conducted most of the experiments and analysis. They used demyelination models to perform *ex vivo* Ca^2+^ imaging, longitudinal *in vivo* Ca^2+^ imaging, *in vivo* MKT-077 infusions, viral injections, CNO activation, immunostainings. L.R.B. conducted rodent and primary human cell culture experiments under the supervision of D.A.M. B.M-S. contributed with widefield Ca^2+^ imaging experiments. C.H. performed the two-photon Ca^2+^ imaging experiments. P.B., D.A.M., B.M-S wrote the FIJI/ImageJ plugin. M.C.A. and F.R. designed the Python software and F.R. wrote the code. M.C.A. performed electrophysiological experiments and data analyses. N.V. performed neurosurgeries and obtained human brain tissue. M.C.A., D.A.M. and R.B-V. wrote the first version of manuscript.

