## Supplementary Figures and Table for "Cell-autonomous mitochondrial calcium flux governs oligodendrocyte regeneration"

**Maas et al.**

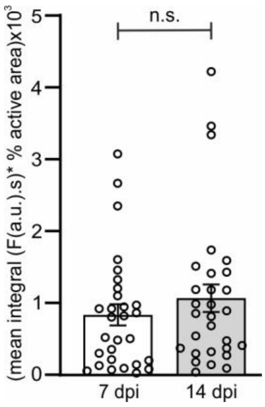


**Supplementary Figure 1 – Oligodendroglial $Ca^{2+}$ signaling in callosal lesions of *Pdgfrα$^{CreERT(+/-)}$;Gcamp6f$^{Lox/Lox}$* mice.** Quantification of oligodendroglial $Ca^{2+}$ imaging in callosal lesions at 7 and 14 dpi in *Pdgfrα$^{CreERT(+/-)}$;Gcamp6f$^{Lox/Lox}$* mice. Data represented as mean ± standard error of the mean (s.e.m.). P-value from two-tailed Mann-Whitney test, n.s. not significant.

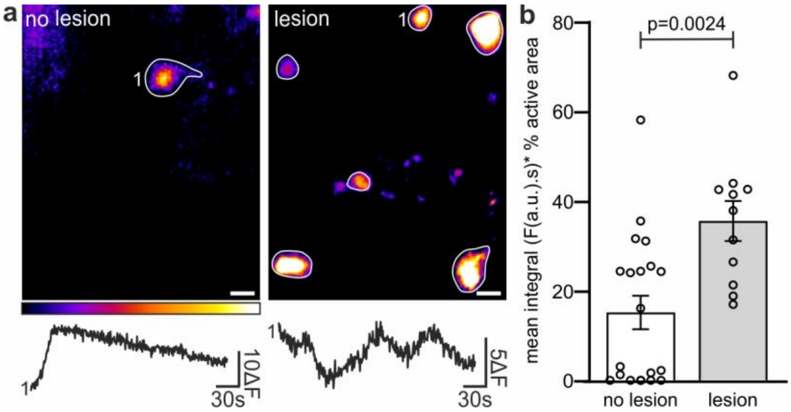


**Supplementary Figure 2 – Demyelination in the corpus callosum activates $Ca^{2+}$ signaling in oligodendroglia.** (**a**) Representative images and $Ca^{2+}$ activity traces of oligodendroglial $Ca^{2+}$ imaging in a normal (no lesion) LPC-induced demyelinated (lesion) corpus callosum of *NG2$^{CreERT2(+/-)}$;GCaMP3f$^{Lox/Lox}$* mice at 7 dpi. (**b**) $Ca^{2+}$ activity is significantly higher in the lesioned than in the non-lesioned corpus callosum Data represented as mean ± s.e.m. p-value from Mann-Whitney test. Scalebar 10 µm.

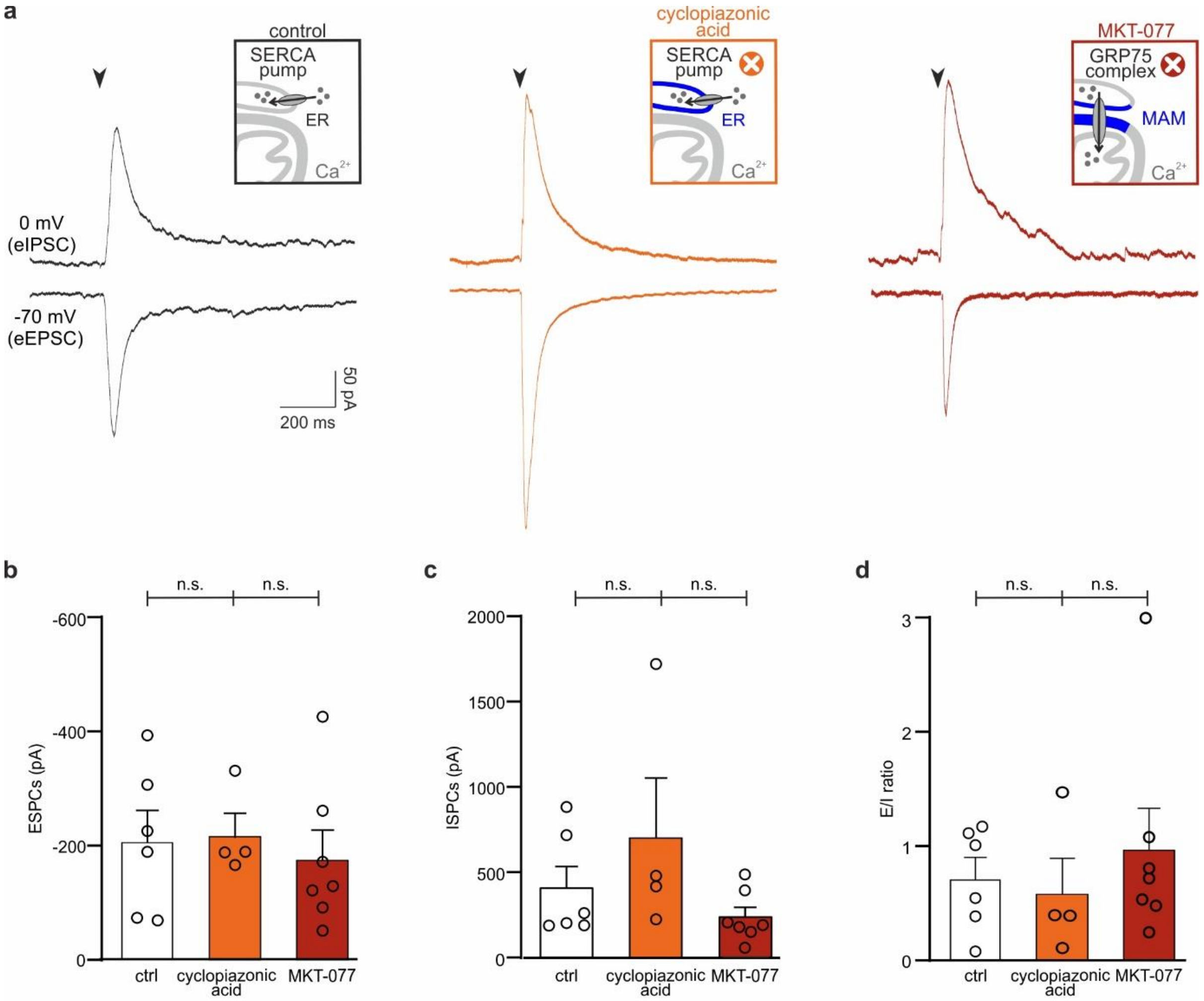


**Supplementary Figure 3 - Inhibition of ER and mitochondrial Ca2+ flux does not alter evoked postsynaptic currents in motor cortex pyramidal neurons.** (**a**) eIPSCs (top trace) and eEPSCs (bottom trace) in layer 5 pyramidal neurons held at 0 mV and −70 mV, respectively, upon extracellular stimulation of acute slices of the motor cortex in control (left) and after more than 30 min incubation with 50 µM cyclopiazonic acid (middle) or after more than 1h incubation with 10 µM MKT-077 (right). Stimulation artifacts blanked, stimulation time indicated (black arrowheads). (**b**) Quantification of eEPSCs (left), eIPSCs (middle) and excitation/inhibition (E/I) ratio (right) calculated from eEPSCs and eIPSCs of layer 5 pyramidal neurons evoked by extracellular stimulation in control (white) and after more than 1h incubation with 50 µM cyclopiazonic acid (orange) or 10 µM MKT-077 (red). Data represented as mean ± s.e.m. n.s.: not significant in Kruskal-Wallis test.

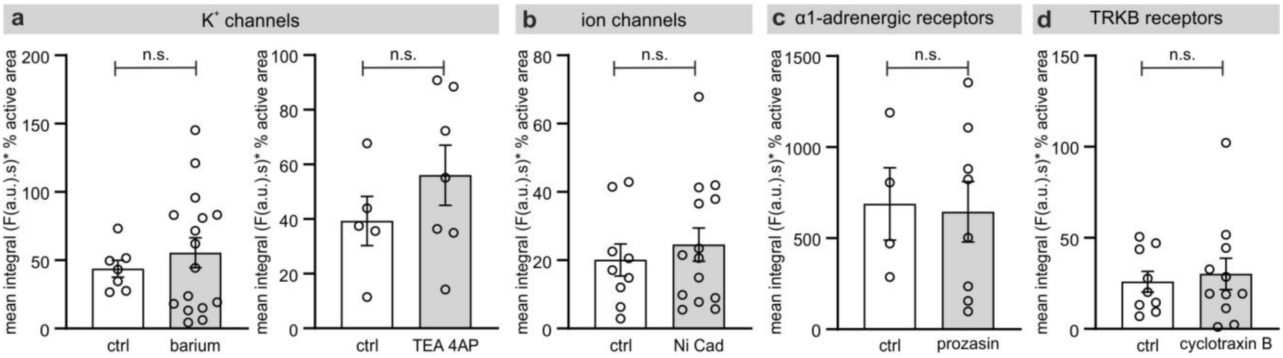


**Supplementary Figure 4 - Quantification of oligodendroglial $Ca^{2+}$ activity in demyelinated lesions following pharmacological blockade.** (**a-d**) Dot plots show quantifications of oligodendroglial $Ca^{2+}$ activity from acute slices at 7dpi under control conditions (white) or in the presence of the $K^+$ blockers Barium or TEA and 4AP (**a**), the voltage-gated $Ca^{2+}$ channel blockers $Ni^{2+}$ and $Cd^{2+}$ (**b**), the α1-adrenergic receptor blocker prozasin (**c**) or the TRKB receptor blocker cyclotraxin B (**d**). P-values from two-tailed unpaired Student's *t* test (**a-c**) and Mann-Whitney test (**d**). Data presented as mean ± s.e.m. n.s. not significant.

**Supplementary Table 1 - Patient characteristics**

| Age | Sex | Diagnosis | Hemisphere | Tissue location |
|---|---|---|---|---|
| 22 | Female | Cavernous haemangioma | Right | temporal |
| 47 | Male | Epilepsy | Right | temporal |
| 53 | Female | Metastatic, breast cancer | Left | unknown |
| 67 | Male | glioblastoma IDH WT, grade IV | Right | temporal |
| 67 | Male | glioblastoma IDH WT, Grade IV | Right | temporal |
| 67 | Female | glioblastoma IDH WT, Grade IV | Right | frontotemporal |
| 75 | Male | Metastasis, non-small cell lung carcinoma | Right | frontal |
