## Supplementary File 1 for "Cell-autonomous mitochondrial calcium flux governs oligodendrocyte regeneration"

#### post-prOccam configuration file for miniscope imaging

[MAIN]

**ACQ_RATE = 0.2**

**FRAME_TOTAL_AREA = 188328**

### The ROIs can be trimmed. To define trimming intervals uncomment the following
### lines and set appropriate values.  The "14 - LAST" format is a convenience
### format to trim the last 14 row of the ROIs. Change the 14 value to any
### appropriate value.

**ROI_TRIM_INTERVALS =**

**1-1449**

**26-LAST**

#175-224

#2 - LAST

### The number of ROI points to be used to compute the
### minimum mean intensity of the ROI.
### The minimum mean intensivy is computed for the following window
### with MIN_MEAN_INTENSITY_HALF_WINDOW = 3:
#
#          [...|...]
#             v
#             min value
### 3 points are selected on the left the of minimum intensity value of the ROI
### and the same number of points are selected on the right of that minimum
### intensity value. The total window thus comprises 7 points.

**MIN_MEAN_INTENSITY_HALF_WINDOW = 20**

```
# The number of ROI points that are used to compute the ROI[z] - ROI[x]
# subtraction over all the ROI intensity values. If MEAN_ABS_DEVIATION_WINDOW =
# 40, the z-x window comprises 40 points.
MEAN_ABS_DEVIATION_WINDOW = 70

# Number by which the mean absolute deviation values are multiplied.
MEAN_ABS_DEVIATION_FACTOR = 5

# Number of points in each ROI that must have an intensity value greater than
# the threshold in order for the ROI to be accepted.
CONDITION_MATCHING_MIN_POINT_COUNT = 4

# Defines the way the trace is "reset" in the ordinate axis.
# Can be one the following values:
#
# - SUB for a simple base line subtraction
# - DFF (F-Fo) / Fo (Delta F / F)
#TRACE_Y_CORRECTION = SUB
TRACE_Y_CORRECTION = DFF

[INTEGRATION]
#
# A list of intervals (unit is the number of the frame
# in the image stack). To set intervals, uncomment the lines below and replace
# with your own values. Remove any unnecessary interval or add any required one.
# After the number interval, put a descriptive label inside double quotes.
# Please, do not comment out the "[INTEGRATION]" line nor the "INTERVALS ="
# line, as the program will fail if these lines are commented out or absent from
# this configuration file.
INTERVALS =
```

**1450-2950**

### 1-2976 "ctrl"

### 155-270 "right after CNO arrives"

### 235-350 "last part of trace CNO for a while"

[CORRELATIONS]

### Set the True if the inter-ROI correlations must be performed.

#PERFORM_ROI_CORRELATIONS = False

PERFORM_ROI_CORRELATIONS = True

### Threshold above which two ROIs are considered correlated.

### This is an absolute value. ROIs will be considered correlated if their

### correlation value is either less than -ROI_CORRELATION_THRESHOLD or greater

### than ROI_CORRELATION_THRESHOLD.

**ROI_CORRELATION_THRESHOLD = 0.9**
